# Structure-Kinetic Relationship for Drug Design Revealed by PLS Model with Retrosynthesis-Based Pre-trained Molecular Representation and Molecular Dynamics Simulation

**DOI:** 10.1101/2022.11.28.518282

**Authors:** Feng Zhou, Shiqiu Yin, Yi Xiao, Zaiyun Lin, Weiqiang Fu, Yingsheng J. Zhang

**Affiliations:** Beijing StoneWise Technology Co Ltd., Haidian Street #15, Haidian District, Beijing 100080, China

## Abstract

Drug design based on their molecular kinetic properties is growing in application. Pre-trained molecular representation based on retrosynthesis prediction model (PMRRP) was trained from 501 inhibitors of 55 proteins and successfully predicted the k_off_ values of 38 inhibitors for HSP90 protein from an independent dataset. Our PMRRP molecular representation outperforms others such as GEM, MPG, and common molecular descriptors from RDKit. Furthermore, we optimized the accelerated molecular dynamics to calculate relative retention times for 128 inhibitors of HSP90. We observed high correlation between the simulated, predicted, and experimental -log(k_off_) scores. Combining machine learning (ML) and molecular dynamics (MD) simulation help design a drug with specific selectivity to the target of interest. Protein-ligand interaction fingerprints (IFPs) derived from accelerated MD further expedite the design of new drugs with the desired kinetic properties. To further validate our k_off_ ML model, from the set of potential HSP90 inhibitors obtained by similarity search of commercial databases, we identified two novel molecules with better predicted k_off_ values and longer simulated retention time than the reference molecules. The IFPs of the novel molecules with the newly discovered interacting residues along the dissociation pathways of HSP90 shed light on the nature of the selectivity of HSP90 protein. We believe the ML model described here is transferable to predict k_off_ of other proteins and enhance the kinetics-based drug design endeavor.

## Introduction

Protein-drug interactions can be described by both thermodynamic properties (K_i_, K_D_, IC50) and kinetic properties (k_on_, k_off_). Ligands with the same affinity may have different association and dissociation rates, so kinetic parameters often provide more useful information for drug design. Compounds having a short retention time in active sites may lead to low drug occupancy at targets and poor efficacy. In contrast, compounds with a longer retention time or slower dissociation rate, their doses can be reduced to achieve better selectivity, and fewer side effects. Studies show that kinetic properties are strongly correlated with the pharmacological activity of drugs in cells and in vivo.^1,2,3,4^ Heitman et al.^2^ studied the relationship among intracellular efficacy, affinity and residence time of ten A_2A_ adenosine receptor agonists, finding that there is a stronger correlation between intracellular efficacy and residence time (R^2^=0.90) than equilibrium constant K_i_ (R^2^=0.13) of these agonists. Because the wet lab experiments to determine the kinetic properties of protein-drug interactions are expensive, new *in silico* methods such as MD simulation and ML are imperative.

Free MD without any bias is the most straightforward way to calculate kinetic properties like k_on_ and k_off_. For example, using the supercomputer ANTON, Shaw et al. firstly reported the binding process of a molecule directly from outside of the pocket by unbiased molecular dynamics.^5^ With 400 μs MD, Pantsar et al. found the important role of water and protein conformational change on the kinetic properties of two p38α MARK inhibitors with short and long residences time but nearly identical activities.^6^ In general, the dissociation time of molecules from their binding pockets are often on the time scale of seconds to hours. Most computing tasks cannot afford such expensive MD simulations. Since 2000, many enhanced sampling methods have been developed to calculate the k_off_. Buch et al.^7^ ran 495 short free MD and used MSM to construct the complete binding process of the trypsin/benzamidine complex, in which 187 trajectories were consistent with the binding mode found in the crystal. Capelli et al. used the infrequent metadynamics (InMetaD) to calculate the k_off_ of the M2 receptor/iperoxo complex, and the result is only an order of magnitude away from the experimental value. InMetaD has also been used to calculate the k_off_ of trypsin-benzamidine^8^, kinase^9,10,11^, biotin-streptavidin^12^, and FKBP^13^. Umbrella sampling (US) was used to study the dissociation process and binding free energy of various protein complexes such as benzamidine-trypsin^14^, acetylcholinesterase^15^, cathepsin K, type I dehydrogenasese, HSP90 and factor Xa. For more complex systems in which the dissociation pathway is straight, the steered MD (SMD) was used together with the US to find the optimal dissociation path and to calculate the potential of mean force (PMF). For example, US+SMD was used to calculate the binding free energy between maltose-binding protein and maltose.^16^ You et al. have applied US+MM/GBSA to study the mechanism of dissociation of two small molecules from ATP allosteric channels of p38 MAP kinase.^17^ SMD was also used to calculate the relative residence times of p38α kinase and FAK ^18,19^. Potterton et al. predicted the relative residence time of 17 small molecules on the A_2A_ adenosine receptor from changes in water-ligand interaction energy.^20^ Mollica et al. used scaled MD (sMD) to calculate the relative residence time of the same series of ligands on proteins such as HSP90, glucose-regulation (Grp78), and the A2A GPCR.^21,22,23^ Zhou et al. combined sMD and InMetaD to calculate the residence time and dissociation mechanism of ASEM.^24^ In 2018, using random accelerated MD (RAMD), Kokh et al. performed large scale k_off_ calculation to calculate the relative residence time of 70 diverse drug-like ligands of N-HSP90 protein and discussed the effect of different substituents on the residence time of small molecules.^25^ The number of inhibitors of N-HSP90 was extended to 94 in a following paper.^26^ They found there is a strong correlation between the calculated retention time (RT) and the experimental value. By combining RAMD and experimental results, Berger et al. found that PF-562271 is more selective for focal adhesion kinase (FAK) kinase than proline-rich tyrosine kinase 2.^27^ RAMD was also used to calculate the dissociation paths of a series of inhibitors of T4 lysozyme, and the results showed disassociation paths of ligands with longer residence time have more intermediate metastable states.^28^ Recently, RAMD was successfully applied to studying the relative residence times, dissociation mechanisms, and the allosteric effects for the two important membrane-embedded drug targets: β2-adrenergic receptor and muscarinic acetylcholine receptor M2.^29^ They found that the dissociation mechanisms observed in the relatively cheap RAMD simulations are consistent with the much more computationally expensive free MD and MetaD simulations. They also uncovered the relationship between the residence time and the allosteric modulation and associated changes in the ligand dissociation pathways. Although there are many successful cases of MD in the calculation of retention time and k^off^, due to the intrinsic drawback of force field and insufficient sampling, the calculated retention time and kzoften deviate from the ground true experimental values especially when the ligands have diverse scaffolds.^25,26,30^ Additionally, most enhanced sampling methods can only produce relative retention times.

In recent years, through the development of artificial intelligence (AI), more accurate predictions of k_off_ were seen by combining molecular simulation and ML. In 2016, a multi-target machine learning (MTML) with interaction fingerprints was used to predict the k_off_ of HIV-1 protease.^31^ Based on electrostatic interaction terms and conformational dynamics, a classification model was used to train and classify 39 inhibitors of HIV-1 protease into four classes, predicting effects with 74% accuracy. This paper argues that electrostatic interactions contribute more to the k_off_ value than van der Waals interactions. Similarly, Zhang et al. used a partial least squares (PLS) regression model to learn interaction features from position-restrained MD to predict the k_off_ of p38 MAPK class II inhibitors and obtained good results.^32^ A 3D QSAR model was also applied to predict k_off_, and Qu et al. used VolSurf descriptors as features to predict the k_off_ of HIV-1 protease inhibitors using PLS regression models.^33^ The Wade group predicted k_off_ with high accuracy using the comparative binding energy (COMBINE) method for HSP90 and HIV-1 protease inhibitors.^34^ The same group also used IFPs from a large number of RAMD trajectories to predict the k_off_ of HSP90.^26^ They found that information along the dissociation pathway is more important than the bound state. Similarly, Huang et al. predicted the k_off_ of 37 inhibitors of HIV-1 using IFPs derived from SMD.^35^. Based on graph methods like contact principal component and pairwise mean Euclidean distance analysis, Bray et al. developed a ML model for clustering the unbinding trajectories and path reaction coordinate detection.^36^ A recent literature reported the prediction of k_off_ for a large array of proteins and ligands using the random forest method with RF-score and protein secondary structures as features.^37^ Although the correlation coefficient r_P_ reaches 0.78, the prediction power of this model became very poor for proteins that had not been seen in the training set. Compared to the traditional molecular simulation methods, the ML method to predict k_off_ still faces many challenges such as proper feature representation and model selection, as well as methods to increase its interpretability to guide drug design.

To enhance ML predicability and increase model interpretability, we trained a new PMRRP to predict the kinetic k_off_ and compared its performance to other popular pre-trained molecular representations (GEM, MPG and other commonly used molecular predictors from RDKit).^38^ Furthermore, by combining the accelerated MD and similarity search, we discovered two molecules having longer retention time for HSP90 than the reference molecules. By combining ML and MD simulation, coupled with the molecular representation features and IFP analysis, we unveil the features that have the most impact on the kinetic properties of the protein target, and the interpretable details on how the molecules achieve their long retention time.

## Materials and methods

### Retrosynthesis-based pretrained representations

We present a novel deep learning model using numerous authentic reaction data for molecular representation learning. Based on our previous work for retrosynthesis prediction [G2GT: Retrosynthesis Prediction with Graph to Graph Attention Neural Network and Self-Training], we cut off the connections between the Encoder and Decoder stage and output the vector of the Decoder as the representation of the input molecule. Because our model utilizes the data of chemical reactions, rather than the traditional single molecule properties, we believe our model extracts more information on molecular functional groups and reactivities, especially certain interactions between molecules that current existing models do not take advantage of during training. Therefore, as a representation that takes consideration of molecular reactivity and intermolecular relationships, our model shows superior performance in property prediction such as k_off_, which involves combination and interaction (complete process of k_off_ prediction in Fig. 1).

**Figure 1.**
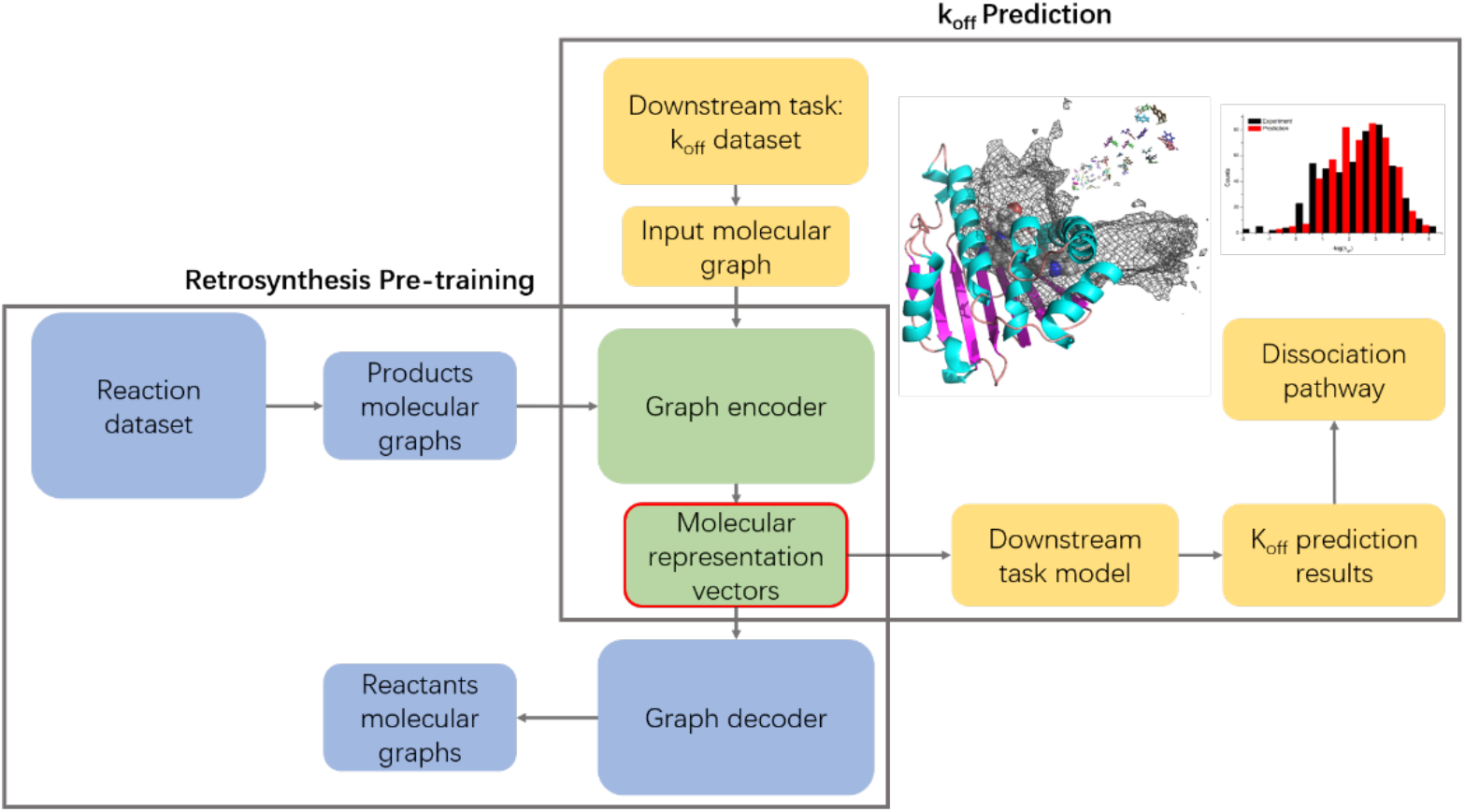
Flowchart of retrosynthesis-based, pretrained representations used for k_off_ prediction

### Partial least squares regression (PLS)

As an extension of linear regression, multiple linear regression (MLR), and principal component regression (PCR), partial least squares regression (PLS) is used to investigate multicollinearity between two groups of variables. For big-p, little-n problems, using PLS to build a model has advantages that MLR and other methods do not have. ^39^ In this paper, we use PLS regression model and PMRRP to predict k_off_.

### RAMD simulations and interaction fingerprints (IFPs)

The relative retention times for the inhibitors were calculated from RAMD, developed by Lüdemann et al.^40^ Short, unbiased MD simulations were independently performed six times for 10ns for each ligand to obtain the equilibrium ensemble of the bound state. A random force was then applied to the ligand to accelerate the dissociation rate of the ligands. The force was set to be 20 kcal/mol/A and the calculated retention time can be controlled to be at a nanoseconds scale. The retention time was calculated as the average value for the 90 independent RAMDs. In all MD simulations, the amber99sb-star-ildn force field ^41^ and TIP3P model ^42^ was used for protein and water. The Gaff2 force field^43^ parameters were generated by ACPYE ^44^ and used for all ligands. The overall temperature of the system was kept constant, pairing independently for each group at 300 K with a V-rescale thermostat. ^45^ The pressure was coupled to a Parrinello-Rahman barostat^46^ at 1 atm separately in every dimension. The temperature and pressure time constants of the coupling were 0.1 and 2 ps respectively. The integration of the equations of motion was performed by using a leapfrog algorithm with a time step of 2 fs. Periodic boundary conditions were implemented in all systems. A cutoff of 1 nm was implemented for the Lennard–Jones and the direct space part of the Ewald sum for Coulombic interactions. The Fourier space part of the Ewald splitting was computed by using the particle-mesh-Ewald method^47^, with a grid length of 0.12 nm on the side and a cubic spline interpolation.

## Results

### k_off_ prediction using pre-trained molecular representation based on retrosynthesis prediction (PMRRP) and cross validation

A common method to measure a pre-trained model’s performance is to test and evaluate it on downstream tasks. ^48^ In order to show that our model can learn a rich and generalized representation, especially for downstream k_off_ tasks, we use different pre-trained models (i.e. MPG and GEM) to generate representation, and then use a unified predictor to predict the k_off_ value. MPG is a pre-trained molecular representation based on molecular graphs.^49^ It combines the classical Neural Message Passing (MPNN) for Quantum Chemistry framework with the powerful transformer block to learn molecular representations. GEM is also a GNN-based molecular pre-training representation.^50^ In order to solve the problem of traditional GNN that three-dimensional molecular information will partly loss in graph representation^51^, GEM introduces molecular graphs and bond angle graphs in which bond-angle information are calculated and shared to jointly model molecular representation. Our PMRRP and the two aforementioned models are all self-supervised pre-training molecular representations. To show the superiority of PMRRP molecular presentation, the traditional RDKID atomic features are also added to our comparison list. The training data is from the 501 inhibitors used by Amangeldiuly et al.^37^ We adopt the leave-one-out strategy to train and predict these samples (Table 1). Our PMRRP molecular representation gives the best prediction results for the k_off_ task, which refers to a higher Pearson correlation coefficient (r_P_= 0.76), compared with GEM (r_P_=0.47) and MPG (r_P_=0.65), and lowest MSE (0.74).

**Table 1.** Comparing different molecular representation architecture in 501 inhibitors -log(k_off_) data, for the prediction stage, we use partial least squares regression (PLS) method.

| | $r_p$ | MSE |
| --- | --- | --- |
| PMRRP | 0.76 | 0.74 |
| RDKit | 0.58 | 1.19 |
| GEM | 0.47 | 1.34 |
| MPG | 0.65 | 1.01 |

### 2. Test of pretrained model on the new database

To demonstrate the generalizability of our pre-trained model, we tested 38 ligands on an independent data set. The 38 ligands are all from the same target protein, HSP90.^57^ HSP90 is a chaperone protein that assists in protein folding, consisting of more kinetic data than other proteins. Furthermore, there are lots of literatures on k_off_ simulation and prediction for HSP90.^52,53,26,54,25,34,55,56,57^ Fig. 2 showed that our prediction value has similar high correlation and lower MSE than the leave-one-out validation. This suggests that our pre-trained molecular representation has superior quality for cross-dataset representations and exhibits high performance in other molecular property prediction task.

**Figure 2.**
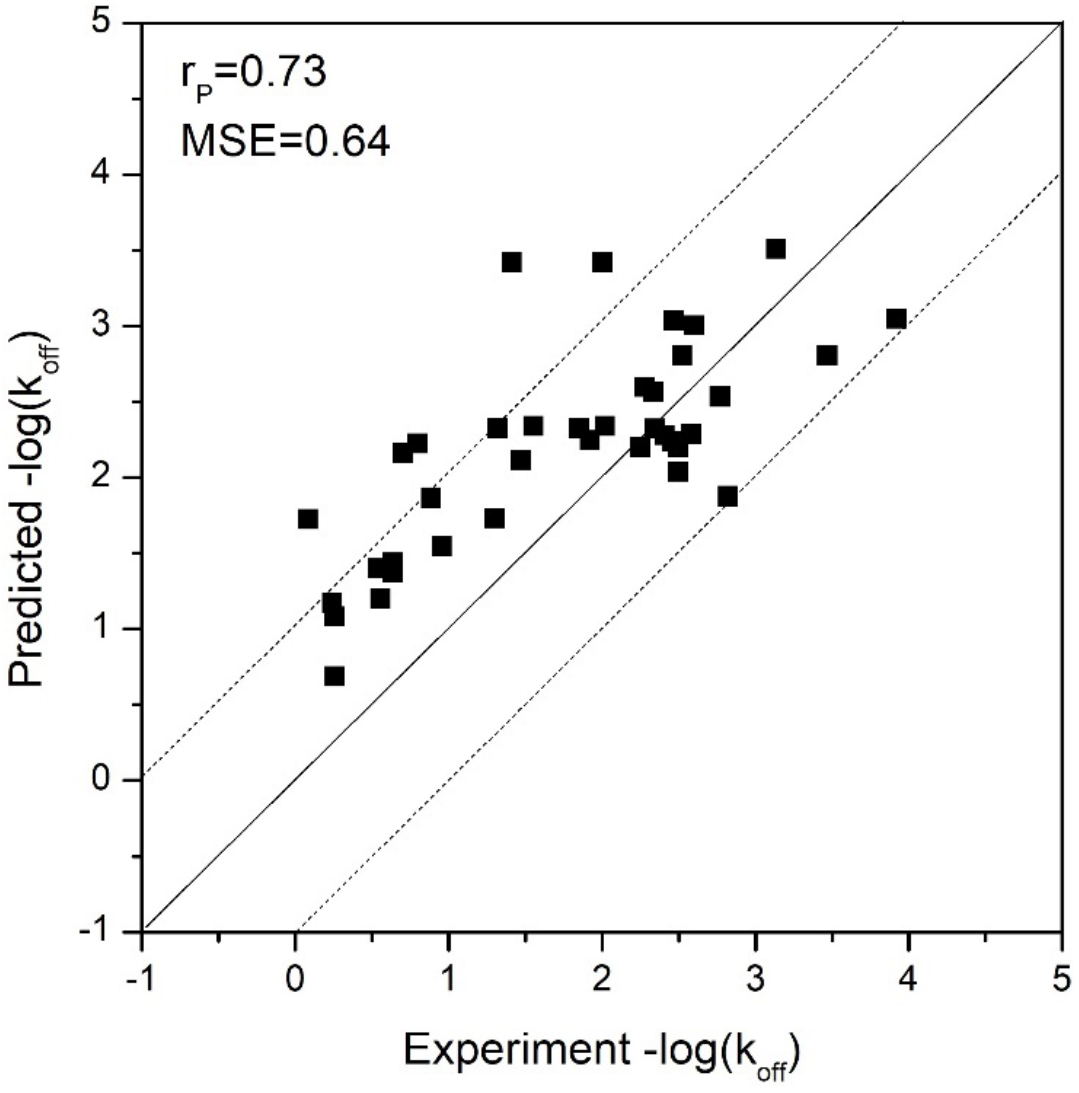
k_off_ prediction on the new 38 inhibitors of N-HSP90 from the independent database.

### 3. Accelerated molecular dynamics unveil the important residues for k_off_ of HSP90 and guide the kinetic-based drug design

The k_off_ prediction by pre-trained molecular representations and PLS model is based only on the molecular features and lack of the protein pocket information. For example, the same molecule may have different binding affinity and k_off_ to different targets. To increase the selectivity of the target, one needs to both consider the features of molecules and the pocket information and protein-ligand interactions. RAMD is a good method to rapidly obtain the relative retention time and dissociation pathways. We performed 90 independent RAMD simulations starting from six different initial structures for each complex to avoid stochastic error. The calculated -log(k_off_) was plotted against experimental values (Fig. 3).

**Figure 3.**
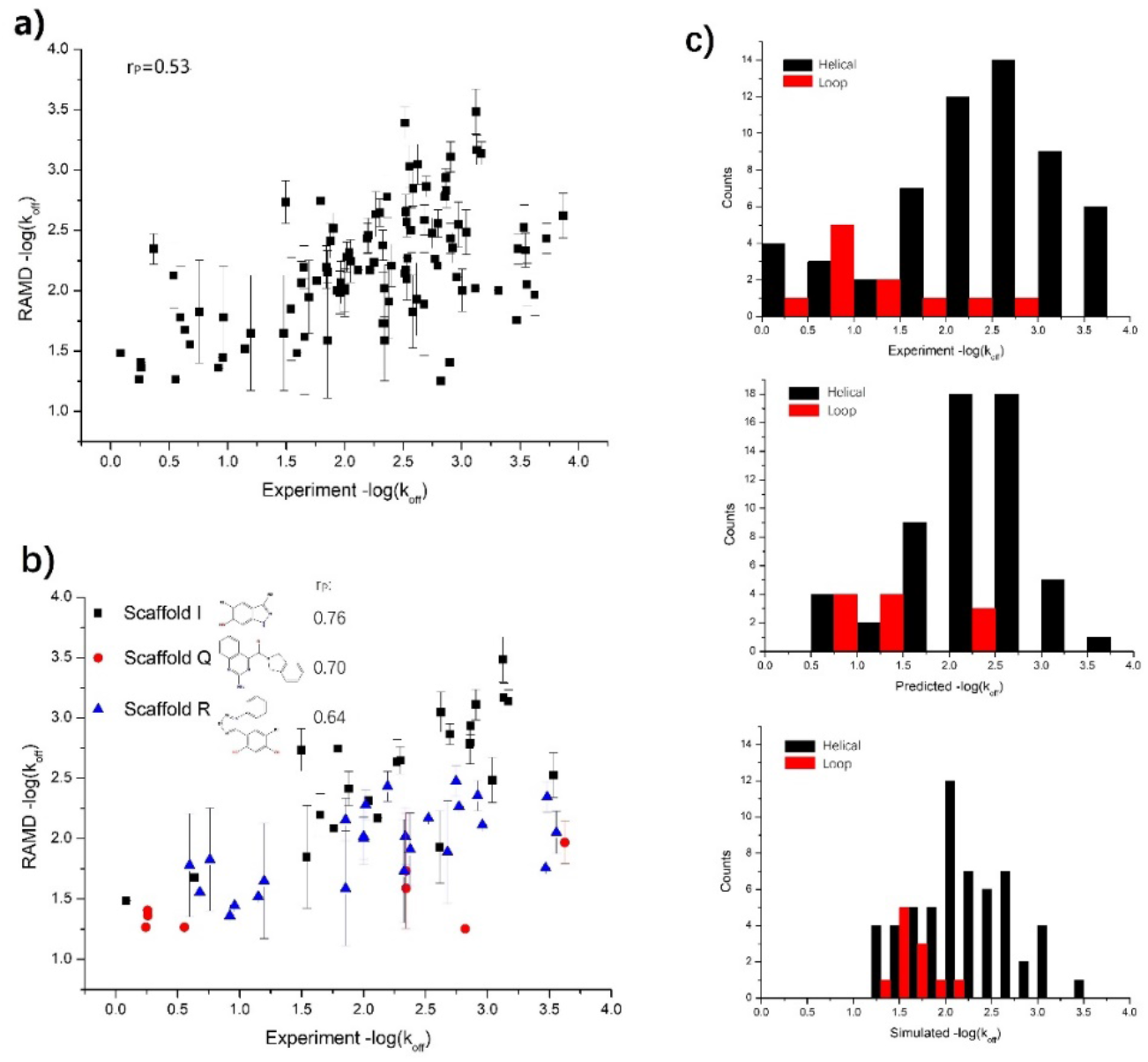
Simulation results for 100 inhibitors of N-HSP90. Scaled residence times plotted vs. experiment -log (k_off_) values on a logarithmic scale for a) the complete 100 sets of compounds. b) grouped by the different scaffolds. Scaffold I = hydroxy-indazole, Scaffold Q = amino-quiza-zoline, Scaffold R =resorcinol. c) The -log(k_off_) population for helical and loop conformation. The correlation coefficient for each set is labeled. The k_off_ is scaled according to the linear fitting -1.2*log(k_off_) + 3 of all compounds.

The Pearson correlation coefficient between the calculated -log(k_off_) from RAMD and the experimental values is not high (r_P_=0.534) for the 100 inhibitors from dataset A. Many outliers, which were also discovered by Kokh et al ^25,26^, were identified from many aspects including the structure of the ligands, the binding mode, and the force field. However, the correlation between the calculated and the measured log(k_off_) is improved when grouping the 100 inhibitors by scaffolds. For example, the calculated Pearson correlation coefficient for scaffold I which contains the hydroxy-indazole structure is 0.76. Kokh et al. also found that the retention time for the 10 compounds of the amino-quinazoline and amino-pyrrolopyrimidine was systematically underestimated.^25^ Consistent with the experiment, we also found that ligands bound to the helical conformation display slow dissociation rates in comparison with those bound to the loop conformation (Fig. 3c).^58^ The predicted and simulated -log(k_off_) value have similar population as the experiment values (i.e. the -log(k_off_) has the largest population in the range from 2.0 to 3.0 for helical conformation). We also performed RAMD simulation on the inhibitors of N-HSP90 protein from Database B. The inhibitors which does not share common scaffold with others are discarded and the remaining 28 new inhibitors were used for RAMD simulation. Both the predicted -log(k_off_) values and simulated ones have high correlation with the experimental values (r_P_=0.73) (Fig. in suppl. info).

Next, we use the IFPs extracted from RAMD as features to study their relationship to the kinetic properties of HSP90 inhibitors. The IFPs were generated from the 90 RAMD trajectories using the python script from Kokh et al.^30^ The IFPs consist of hydrogen bond donor (HD), hydrogen bond acceptor (HA), hydrophobic (HY), salt bridges (IN), π-π stacking (AR), and Halogen-π interaction (HL).

From the PLS coefficient (Fig. 4), one can analyze the most important IFPs which have a strong correlation with the kinetic properties of ligands. These significant IFPs involve residues like ASN51, SER52, ASP54, ASP93, GLY97, ASP102, and LEU107, most of which polar and surrounding the ATP binding pocket, TYR139 and THR184, inside the hydrophobic sub-pocket, and PHE138 at the entrance of the hydrophobic sub-pocket. The results are in line with the findings by Kokh et al.^26^ A distal residue ILE110 at the exit of the hydrophobic sub-pocket does not interacts with ligands in the bound state, but shows the importance during the dissociation process. Fig. 5 outlines these important residues and their positions on N-HSP90.

**Figure 4.**
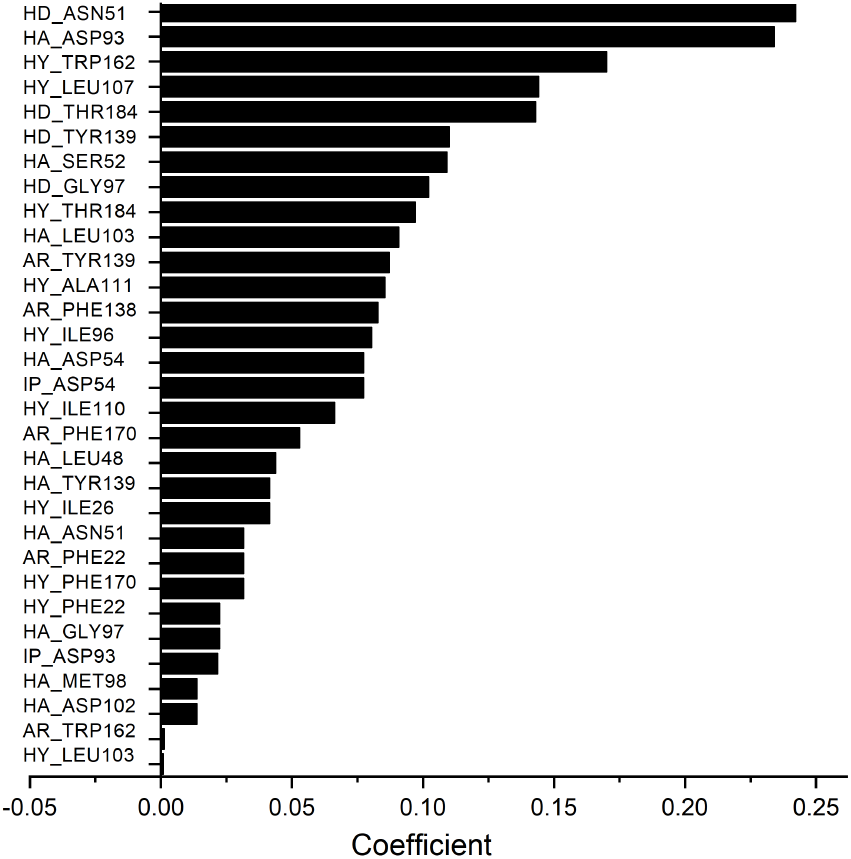
Coefficients of the PLS model in the leave-one-out cross validation built on RAMD IFPs only (only the top 30 are shown).

**Figure 5.**
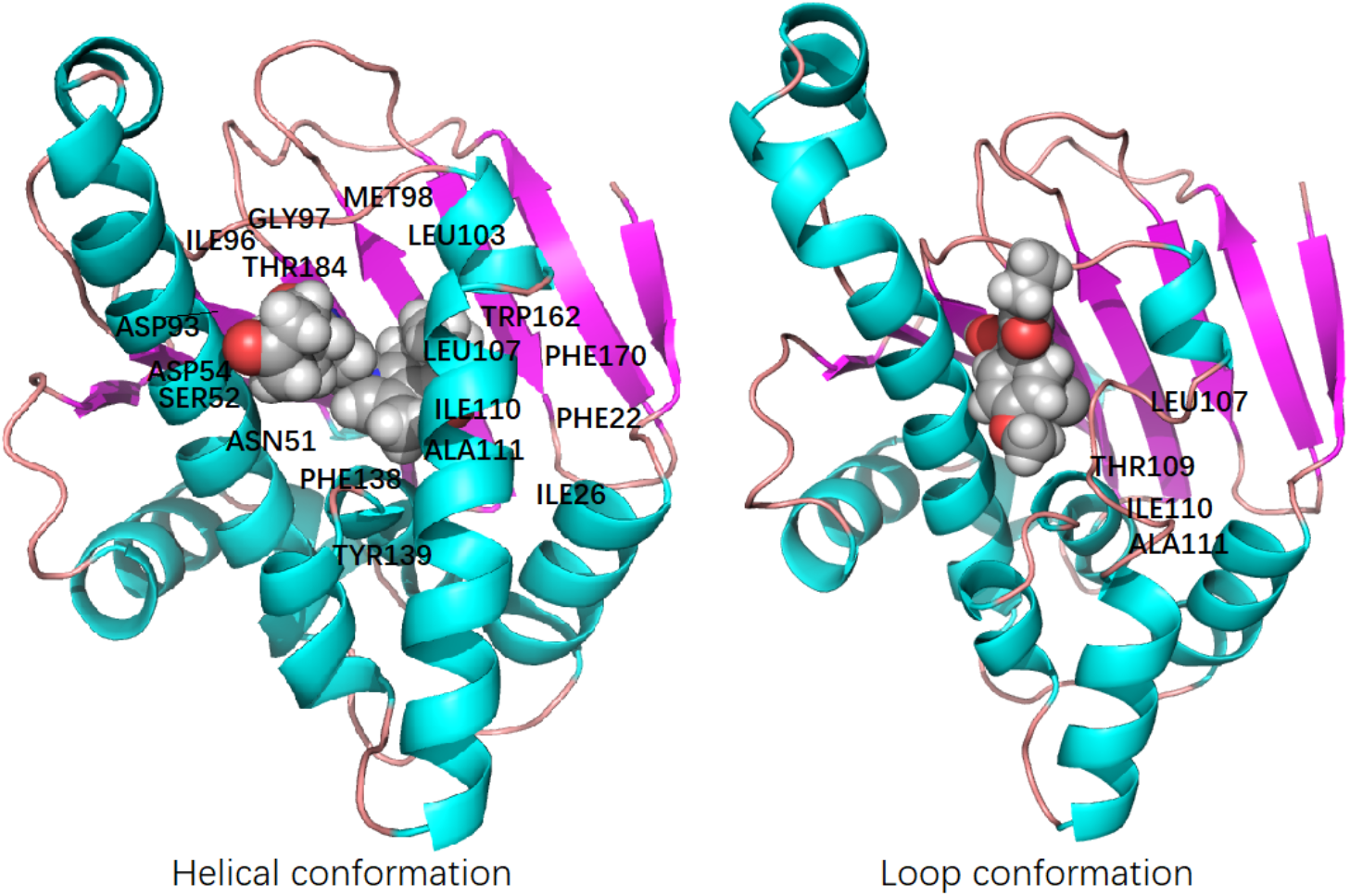
The helical and loop conformation of HSP90, the important residues are labeled.

Fig. 6 shows the schematic visualization of the RAMD dissociation trajectories of the four compounds with slowest (Fig. 6a, b) and fastest (Fig. 6c, d) dissociation rate among the simulated 28 inhibitors from Database B. There are much more interactions of 5j6m_ligand than 6ei5_ligand along the dissociation pathways. Aforementioned important residues, ASN51, ASP93, GLY97, PHE138, and THR184, are all existent and retained during the dissociation process for 5j6m_ligand. However, for 6ei5_ligand, these interactions are either absent or transient during the dissociation process. Furthermore, the dissociation pathways show that for 6ei5_ligand the transition flow between each intermediate is small while the transition to the unbound state has a large flow, which suggests a fast dissociation. On the other hand, for 5j6m_ligand there is a large transition flow between Clusters 1 to 6 before it transits into Cluster 7 which is the last intermediate before fully unbinding. That means it costs more time for 5j6m_ligand to transit between different intermediates, and the transition barrier to the unbound state is higher.

**Figure 6.**
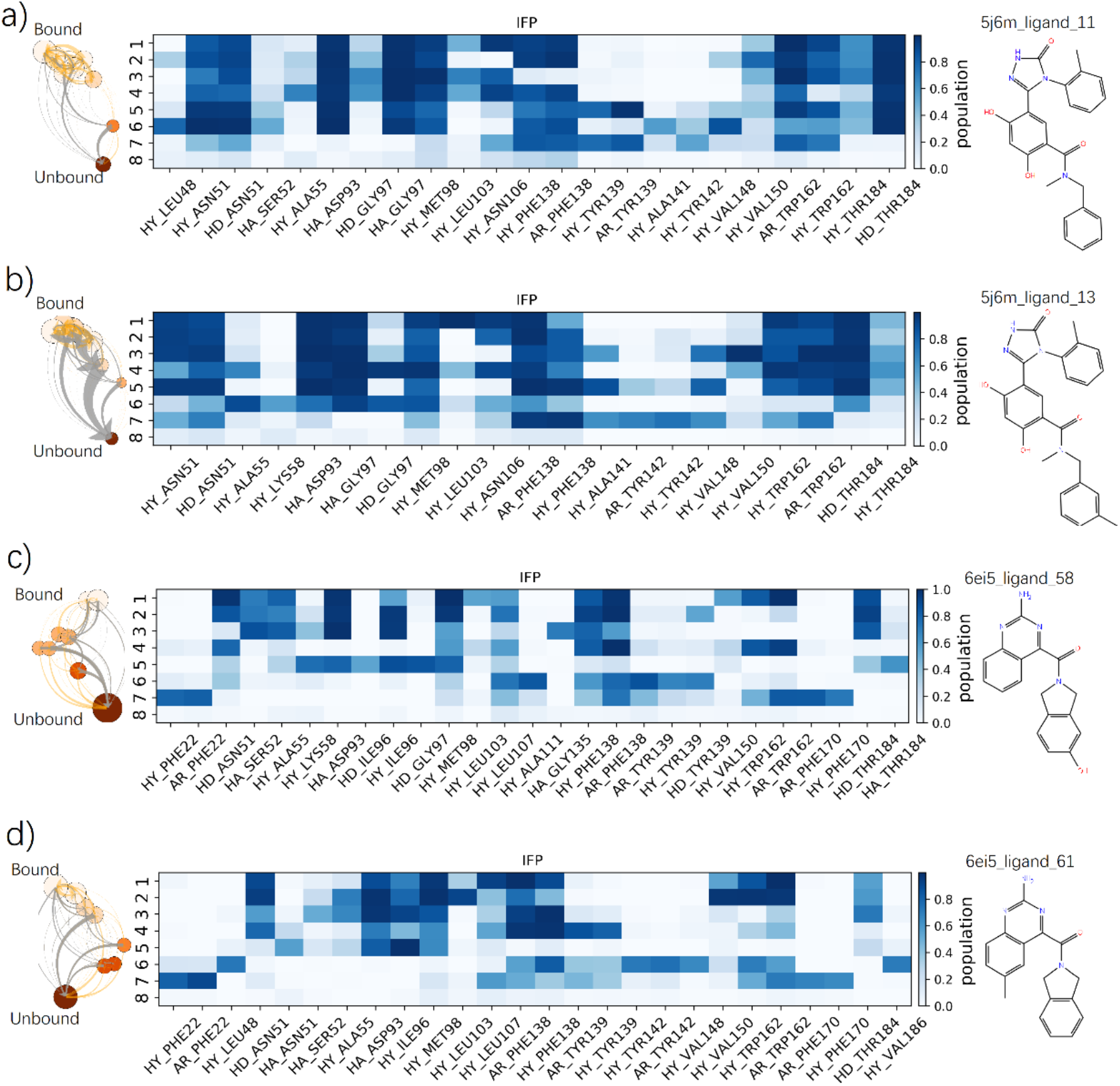
Schematic visualization of the RAMD dissociation trajectories of 5j6m_ligand and 6ei5_ligand of N-HSP90. From the left to right shows: 2D graph representation of the dissociation pathways; IFP composition of each cluster along the dissociation pathways (Cluster 1 is the bound state, and Cluster 8 is the unbound state); Chemical structures of the compound. Each cluster is shown by a node with the size indicating the cluster population. The light color is the bound state and the dark color is the unbound state. The width of the light-orange arrows is proportional to the number of corresponding transitions between two nodes, and the gray arrows indicate the total flow between two nodes.

**Figure 7.**
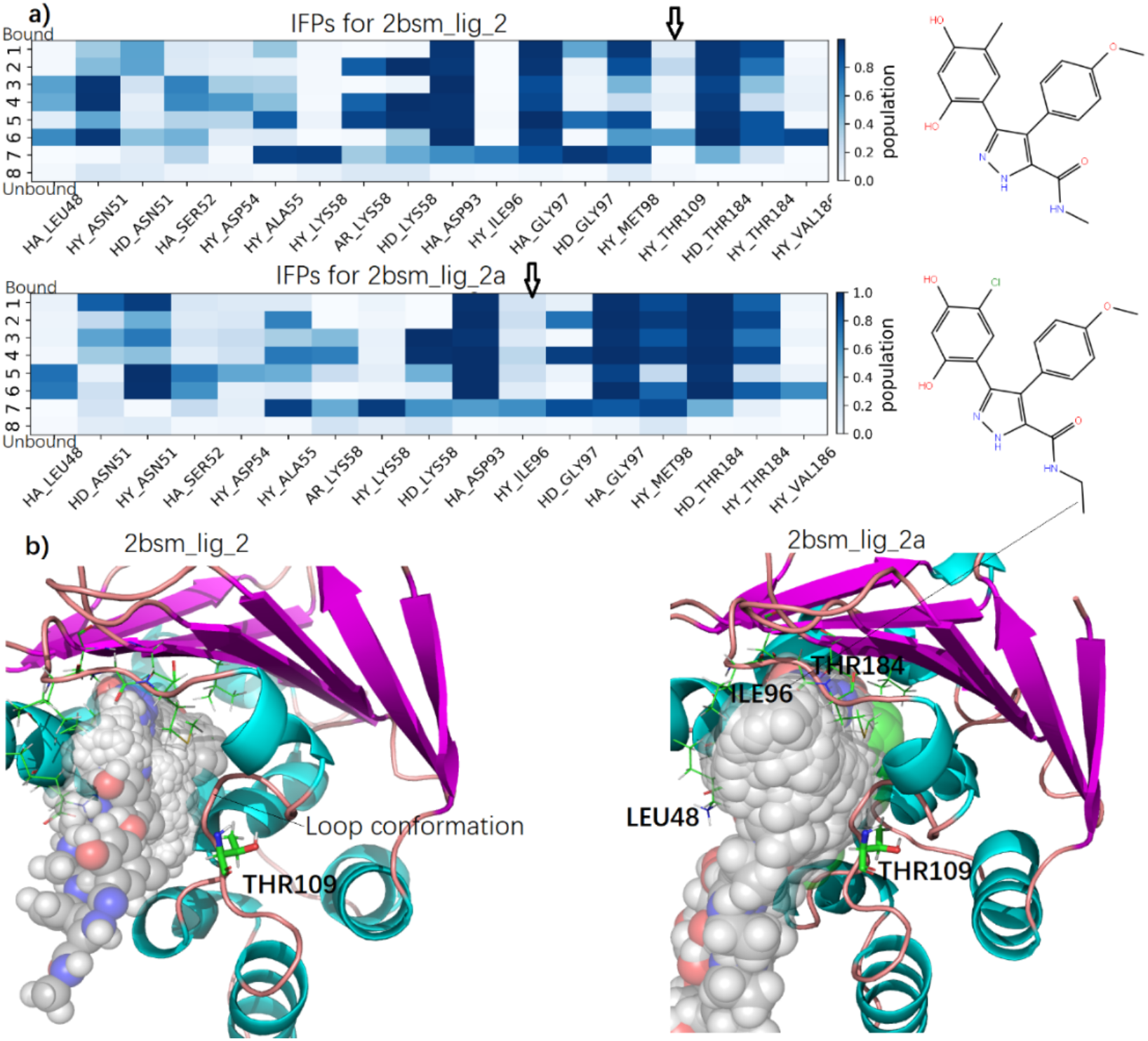
Schematic visualization of the RAMD dissociation trajectories of 2bsm_lig_2 and 2b-sm_lig_2a. a) IFP composition of each cluster along the dissociation pathways and the chemical structures of the compound. b) The pocket and the important residues along the dissociation pathway for 2bsm_lig_2 and 2bsm_lig_2a. The ligand is show in sphere for each frame along the dissociation pathway.

**Figure 8.**
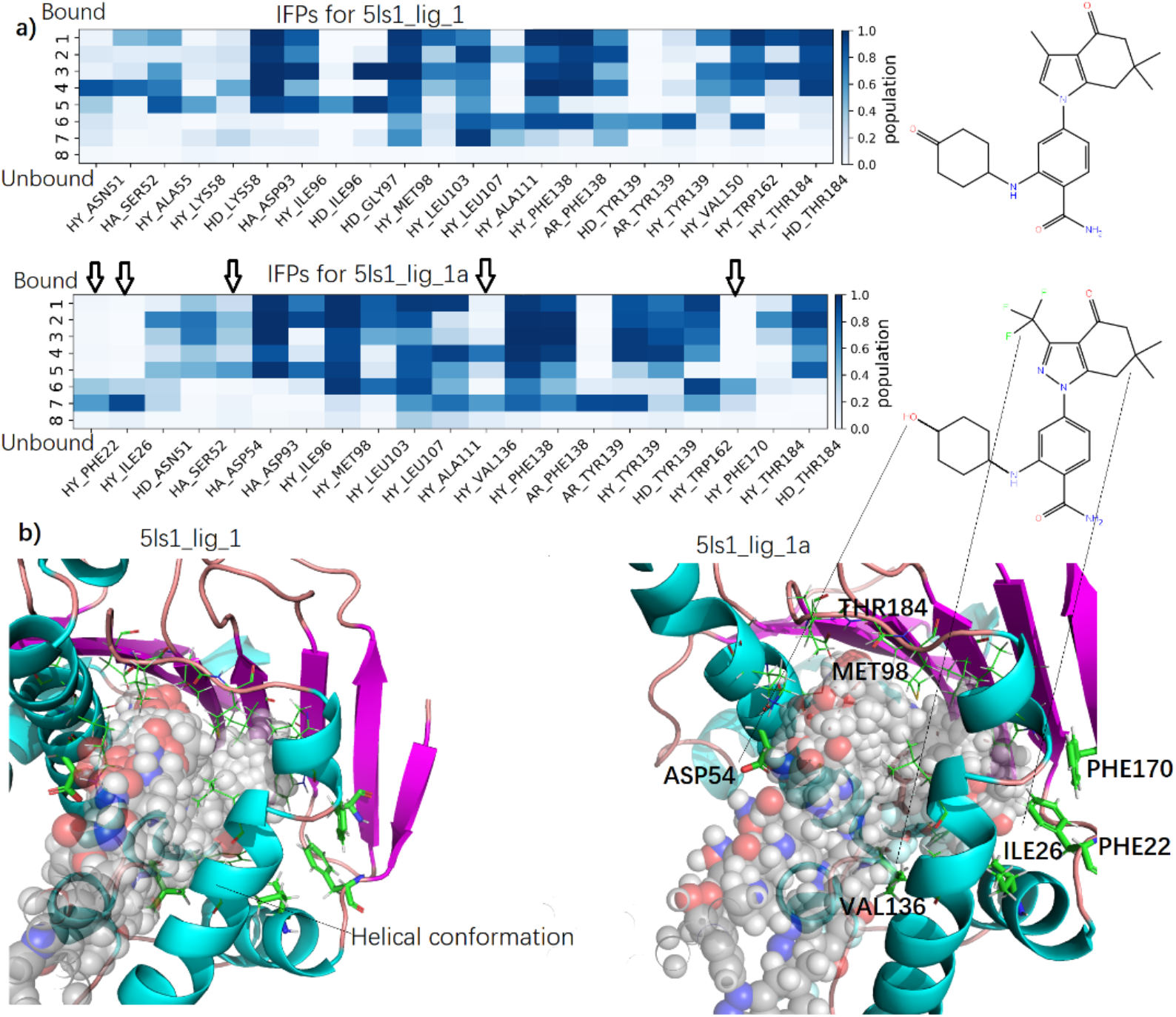
Schematic visualization of the RAMD dissociation trajectories of 5ls1_lig_1 and 5l-s1_lig_1a. a) IFP composition of each cluster along the dissociation pathways and the chemical structures of the compound. b) The pocket and the important residues along the dissociation pathway for 2bsm_lig_2 and 2bsm_lig_2a. The ligand is shown in sphere for each frame along the dissociation pathway.

### 4. New molecules from similarity search - A case study of structure-kinetic relationship for drug design

One strategy to increase selectivity of drugs is to optimize the kinetic properties in order to obtain the drug with longer retention time. Based on kinetics-based drug design, Khanna et al. optimized the kinetic property of EZH2 Inhibitors and achieved 10 times increase of retention time only by changing -OCH_3_ to -SCH_3_.^59^ Zhou et al. designed and synthesized 44 novel derivatives containing an important fluorine atom, with much lower k_off_ than donepezil.^60^ Among them, compound **12** demonstrated a much better efficacy and a lower effective dose than that of donepezil. By using ensemble molecular docking and relative binding free-energy calculations Miller et al. found a novel compound TDI-11861 which shows both improved sAC (ADCY10) binding affinity and residence time than its precursor (3181 s versus to 25 s).^61^ Combining induced fit docking and MM/GBSA calculation cubic by cubic, Bai et al., construct a sophisticated energy landscape of ligand dissociation and found a potential lead compound which shows better binding kinetic and thermodynamic properties with either TcAChE or mAChE.^62^ Here, we combine ML, accelerated MD and similarity search to find potential molecules with longer retention time than their precursors. The databases used for similarity search includes Taosu, MCE, Enamine and Selleck. We applied docking and modeling to guarantee the ligands we identified have the similar binding modes. To demonstrate the approach’s generality and novelty of identified molecules, we only consider the molecules with similarity score lower than 0.35. Ten molecules are discovered from this process. Among them, two are identical reference molecules and discarded. The k_off_ values are predicted for the rest eight novel molecules, two of them have improved k_off_ than their references (Table 2).

**Table 2.** The similarity score and the predicted -log(k_off_) for reference compounds and searched compounds.

| Reference compound name | Similarity score | Predicted $-\log(k_{\text{off}})$ of template compound | Predicted $-\log(k_{\text{off}})$ of searched compound |
| --- | --- | --- | --- |
| 2uwd_lig_1 | 0 | 2.40 | 2.40 |
| 6eln_lig_1 | 0 | 1.98 | 1.98 |
| **2bsm_lig_2** | **0.19** | **2.06** | **2.30** |
| 2bsm_lig_4 | 0.2 | 2.44 | 2.30 |
| 6f1n_lig_1 | 0.22 | 1.89 | 1.22 |
| 2bsm_lig_1 | 0.25 | 2.20 | 1.79 |
| 2bsm_lig_3 | 0.26 | 2.35 | 1.77 |
| **5ls1_lig_1** | **0.33** | **3.26** | **3.42** |
| 5j64_lig_2 | 0.33 | 1.97 | 1.77 |
| 5j64_lig_4 | 0.34 | 2.33 | 1.77 |

We use accelerated MD to calculate the relative retention time for the two discovered new molecules (2bsm_lig_2a and 5ls1_lig_1a). The simulated -log(k_off_) is 1.56 and 1.61 for 2bsm_lig_2 and 2bsm_lig_2a, and 1.58 and 1.69 for 5ls1_lig_1 and 5ls1_lig1a. Both prediction and simulation show improved k_off_ for 2bsm_lig_2a and 5ls1_lig_1a over their reference molecules. To understand the mechanism behind the improved kinetic properties, we analyzed the trajectories and plot the IFPs along the dissociation path.

First, we compared the chemical structure and dissociation pathway of 2bsm_lig_2 and 2b-sm_lig_2a. 2bsm_lig_2 is an inhibitor for the loop conformation of HSP90. Compared to 2b-sm_lig_2, 2bsm_lig_2a has stronger interactions with ILE96 due to the longer aliphatic chain of ethyl group than methyl group. However, there is weak hydrophobic interaction with THR109 for 2bsm_lig_2, which is absent for 2bsm_lig_2a. From Fig 4, we note that THR109 is not the important residue for the k_off_ value, so we argue the interaction between 2bsm_lig_2 and THR109 is trivial. The role of changing from methyl group to chlorine group is unclear, a smaller sized chlorine group may help alleviate some strain energy when the -CH_3_ grow to -CH_2_CH_3_.

Next, we compare the chemical structures and dissociation pathways of 5ls1_lig_1 and 5l-s1_lig_1a. 5ls1_lig_1 is an inhibitor for the helical conformation of HSP90. There are two structural differences between the two molecules. First, 5ls1_lig_1a has -CF_3_ instead of -CH_3_, which has extra interaction with VAL136. Since VAL136 is at the entrance of the binding pocket, the larger -CF_3_ group may have stronger interaction with isopropyl group of VAL136 than -CH_3_, thus decreasing the dissociation rate. The other pronounced change is from carbonyl group to hydroxyl group, which leads to a H-Bond to ASP54. ASP54 is also at the entrance, the newly formed H-Bond may also reduce the dissociation rate. PHE22, ILE26, and PHE170, which are all important residues as show in Fig. 4, form interactions just before the ligand fully dissociated from the protein, indicating a potential intermediate structure. After checking the trajectories, we found one configuration in which the -CF_3_ and two -CH_3_ groups on the six-membered ring have hydrophobic interactions with PHE22, ILE26 and PHE170 (Fig. 9). This intermediate structure has short retention time and quickly dissociates.

**Figure 9.**
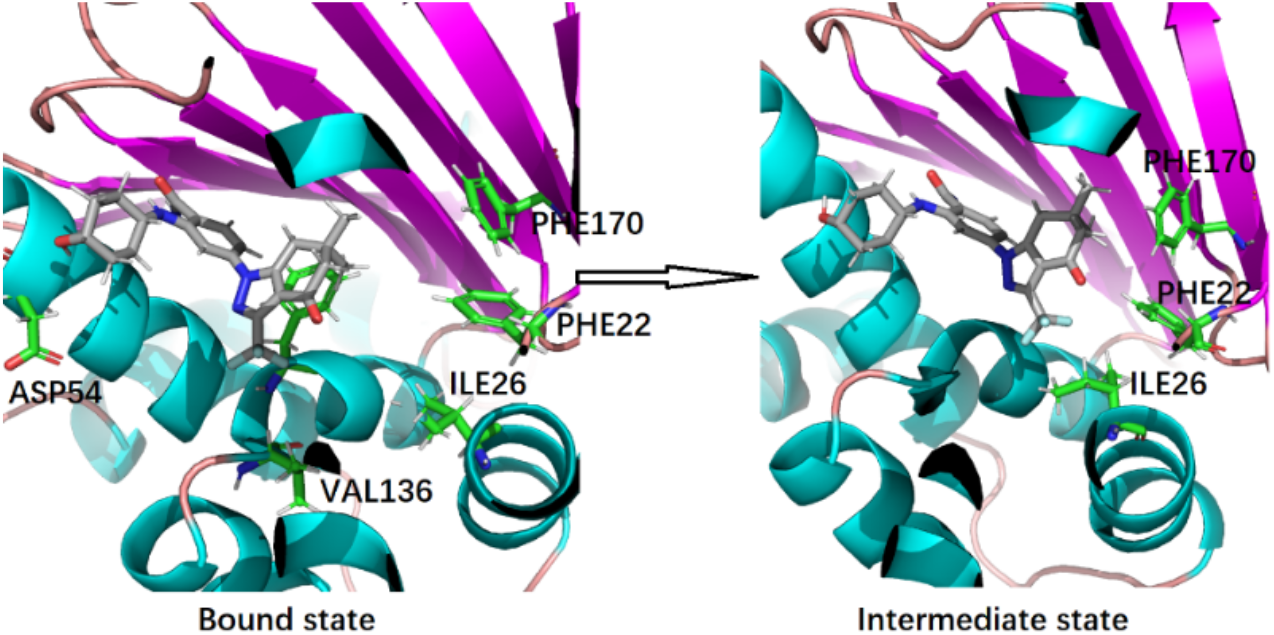
The pocket and the important residues for the bound state and intermediate state of 5l-s1_lig_1a bound complex.

For these promising molecules, we can further use more accurate simulation methods such as free energy perturbation or umbrella sampling to validate the binding affinity before engaging in wet lab testing.

## Conclusion

The PMRRP-based PLS ML model was successfully applied to predict the k_off_ of 138 inhibitors of N-HSP90 protein. The relative retention time is calculated using RAMD simulation. Both methods can produce the -log(k_off_) which has high correlation to the corresponding experimental data. The molecular features from the pretrained model and the IFPs from RAMD together give more information for the drug design. Using our ML k_off_ prediction approach, we discovered two novel compounds, [2bsm_lig_2a and 5ls1_lig_1a], from a set of potential hits obtained using similarity search, and demonstrated the improved retention times for N-HSP90 protein. The resulting molecular mechanisms were elucidated both in molecular features and IFPs along the dissociation pathway. Our proposed protocol offers a feasible way for kinetics-driven drug design and increases the success of finding molecules with proper kinetic profiles to the interested targets.

## Author

Feng Zhou - Beijing StoneWise Technology Co Ltd, Haidian street #15, Haidian district, Beijing, Beijing, CN 100097

Shiqiu Yin - Beijing StoneWise Technology Co Ltd, Haidian street #15, Haidian district, Beijing, Beijing, CN 100097

Yi Xiao - Beijing StoneWise Technology Co Ltd, Haidian street #15, Haidian district, Beijing, Beijing, CN 100097

Zaiyun Lin - Beijing StoneWise Technology Co Ltd, Haidian street #15, Haidian district, Bei- jing, Beijing, CN 100097

Weiqiang Fu - Beijing StoneWise Technology Co Ltd, Haidian street #15, Haidian district, Bei- jing, Beijing, CN 100097

## Notes

The authors declare no competing financial interest.

## ACKNOWLEDGMENT

We acknowledge and thank Jielong Zhou, Mengcheng Pu, Biao Fu, and Yang Wang for helpful discussions and comments on both the method and this paper.

